# Intracortical microstructure profiling: a versatile method for indexing cortical lamination

**DOI:** 10.1101/2025.10.16.682836

**Authors:** Casey Paquola, Jessica Royer, Thanos Tsigaras, Donna Gift Cabalo, Youngeun Hwang, Felix Hoffstaedter, Simon B Eickhoff, Boris C Bernhardt

## Abstract

Intracortical microstructure profiling represents a powerful, scalable approach for investigating the laminar organisation of the human cortex on both *in vivo* and *post-mortem* datasets. Building upon a long tradition of histological analysis, this method leverages surface-based intracortical sampling to generate profiles of tissue properties across cortical depths. The present work outlines a standardised workflow for intracortical microstructural profiling, newly packaged as an open-source toolbox “CortPro” (https://github.com/caseypaquola/cortpro). Here, we explore the utility of central moments as descriptors of profile shape. Using these measures, we quantify *(i)* the extent to which in vivo MRI can capture laminar differentiation, *(ii)* the test-retest reliability of profiles, and *(iii)* their replicability across sites and studies. Our results demonstrate that intracortical profiles are remarkably robust and effectively mitigate bias-field related limitations of non-quantitative MRI. As applications of microstructure-sensitive imaging expand across development, aging, and disease, microstructure profiling provides a principled means of linking microstructural neuroanatomy with systems-level brain organisation.

## Background and introduction

Intracortical microstructure profiling was first developed in the 1960’s as a method for appraising cortical microarchitecture. It emerged in a period of scepticism about the reproducibility of foundational neuroanatomical maps, owing to their dependence on visual evaluation (Lashley and Clark, 1946; Bailey and von Bonin, 1951; Braitenberg, 1962). Researchers today are similarly concerned about reproducibility, especially in the context of multi-site studies and biomarker development. Here, we aim to demonstrate that intracortical microstructure profiling presents a promising way forward for objective characterisation of cortical microarchitecture, relevant to understanding differences across regions and individuals.

In recent years, intracortical microstructure profiling has been reinvigorated by parallel advances in histology and neuroimaging. On the one hand, new alignment techniques enabled reconstruction of histological sections into 3D volumes (Amunts *et al*., 2013; Alkemade *et al*., 2022), thereby allowing cortical layers to be visualised across the entire human brain regardless of the cutting plane (Novek *et al*., 2023). On the other hand, sub-millimetre, microstructure-sensitive magnetic resonance imaging (MRI) has recently been made feasible, owing to higher magnetic field strengths, acceleration factors and sequencing innovations (Bock *et al*., 2013; Dinse *et al*., 2015; Weiskopf *et al*., 2021).

While microstructure profiling was originally developed for *post-mortem* histology, its translation to *in-vivo* imaging paved the way for a bountiful new era of investigations into cortical microstructure. Recently, researchers have identified differences in cortical microstructure depending on biological (Sprooten *et al*., 2019; Wei *et al*., 2022; Royer *et al*., 2023; Küchenhoff *et al*., 2024; Park *et al*., 2024) or cognitive-behavioural factors (Patel *et al*., 2023; Valk *et al*., 2023; Lee *et al*., 2025), including particularly strong changes with developmental and lifespan processes (Whitaker *et al*., 2016; Paquola, Bethlehem, *et al*., 2019; Sui *et al*., 2022; Hettwer *et al*., 2024; Sydnor *et al*., 2025). More generally, microstructure profiles enabled the first quantitative evidence that the long-discussed sensory-fugal axis predominates cortical organisation (Sanides, 1962; Mesulam, 1998). Furthermore, as the sensory-fugal axis reflects differences in supra-vs infra-granular microstructure, positioning activation patterns or group differences on the sensory-fugal axis can shed light on their position in cortical hierarchies. While this field is still in its infancy, *in-vivo* profiling alongside fMRI avails a huge realm of possibilities for uncovering the dependences between microstructure and function in the human brain and potentially unravelling mechanisms of cognition (Paquola *et al*., 2022; DeKraker *et al*., 2025).

### The Intracortical Microstructure Profiling Workflow

A central methodological consideration in intracortical profiling concerns how sampling points are distributed across cortical depth. Cortical layers vary in thickness as a function of cortical curvature, with upper layers tending to be thicker in sulci and deeper layers thicker in gyri (Bok, 1929). To account for this, sampling points can be positioned using an equivolumetric algorithm, which maintains a constant volume fraction for each cortical segment (Waehnert *et al*., 2014). In essence, this approach ensures that a given fraction of cortical volume is represented equally across regions, much like how the top centimetre of a wide martini glass can contain the same amount of liquid as several centimetres near its narrow base. Operationally, equivolumetric surface generation involves adjusting the depth of each vertex on the intracortical surface mesh according to its position within sulci or gyri. This yields surfaces that more closely follow true cortical lamination compared with alternative approaches such as equidistant or Laplacian-based sampling (Waehnert *et al*., 2014). It is important to note, however, that layer thickness also varies between cortical areas (Von Economo and Koskinas, 1925). Consequently, equivolumetric sampling preserves relative layering within a given area, but the same fractional depth may not correspond to an identical histological layer across areas. Intracortical microstructure profiling is therefore sensitive to patterns of lamination rather than absolute differences within specific layers. This representation nonetheless provides valuable insight into regional microarchitecture and fine-grained cortical differentiation (García-Cabezas, Hacker and Zikopoulos, 2020; Paquola *et al*., 2025).

Secondly, the appropriate density of sampling is dependent on the resolution of the underlying image. Take for example an MRI of an adult brain with isotropic resolution of 0.8mm. Given an average cortical thickness of 2.8mm, one may assume that a profile captures approximately 3.2 unique voxels. In practice the number is significantly higher, because cortical surfaces are variably positioned relative to voxels (mean±SD: 5.6 ± 1.5 voxels, **Supplementary Figure 1**). Additionally, this calculation assumes nearest neighbour interpolation, but trilinear interpolation is typically advised to avoid stair-step artefacts. Trilinear interpolation involves taking a weighted average of eight neighbouring voxels for each sampling point, thereby producing a smooth estimate of continuous transitions in image intensities. Using this technique, the number of unique values per profile drastically increases (mean±SD: 38.7±6.4). In our standard workflow, we recommend striking a balance between these nearest neighbour and trilinear estimates of unique values (*e.g*., 14 samples), thereby limiting redundancy in the profile while still capturing a full spectrum of microstructural variations across cortical depths.

Building on these insights, we developed a standardized workflow for extracting microstructure profiles (MPs) (Paquola, Vos De Wael, *et al*., 2019). The procedure involves: *(i)* reconstructing the cortical surface, *(ii)* generating equivolumetric intracortical surfaces with matched vertex indices, *(iii)* registering a microstructure-sensitive image to surface space, (iv) sampling voxel intensities along the matched intracortical surfaces, and (v) compiling these intensity values into profiles that capture depth-wise variations in tissue properties (**Figure 1**, see **Supplementary Material** for full protocol).

**Figure 1.**
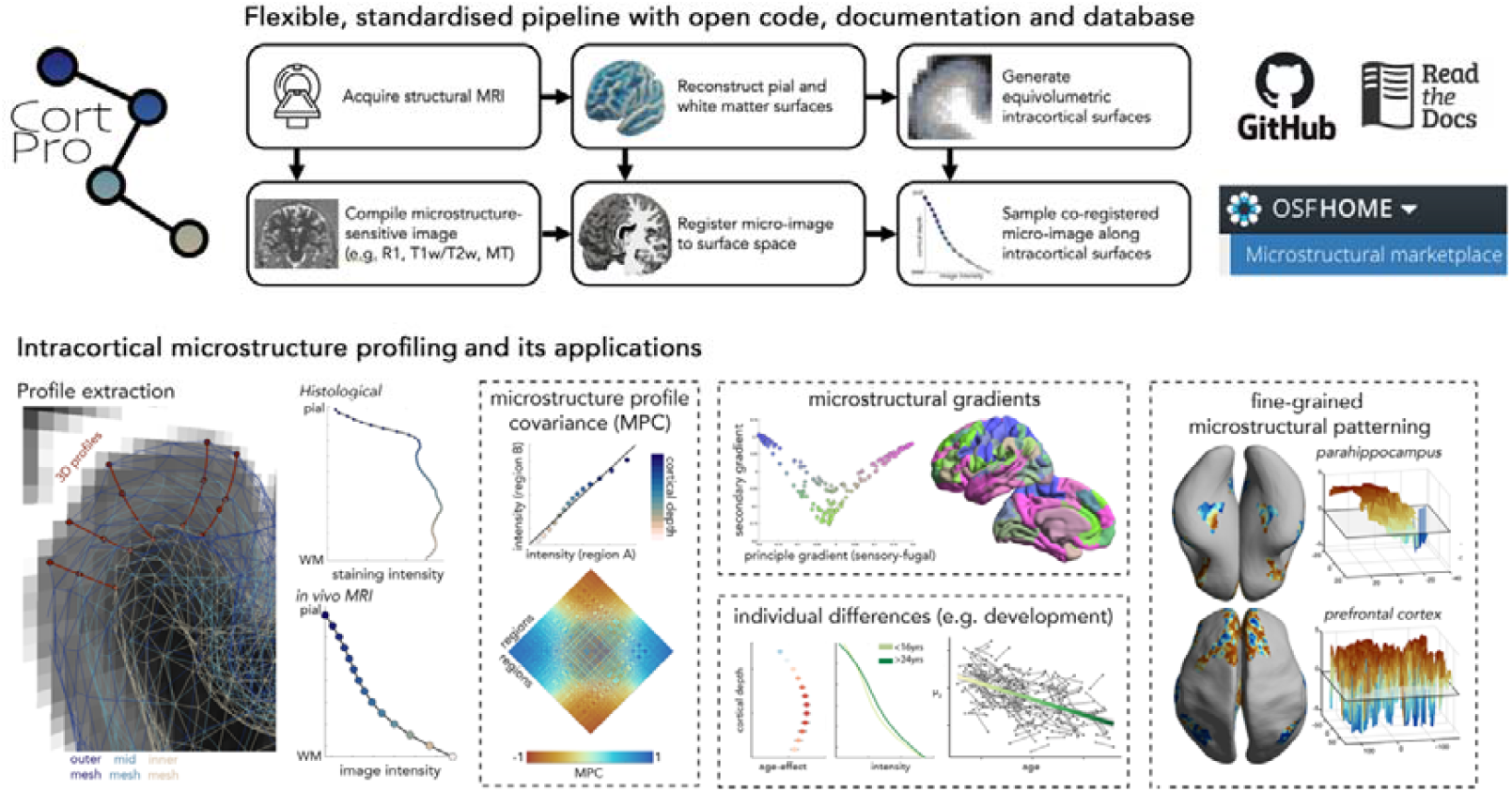
**(Above)** Flowchart of intracortical microstructure profiling approach. The acquisition protocol should include T1-weighted imaging (e.g. MPRAGE) and a high-resolution (≤1mm) sequence that enables a microstructural contrast (e.g. MP2RAGE (Marques *et al*., 2010), SPACE for T2-weighted (Mugler *et al*., 2000) or MPM (Weiskopf *et al*., 2013)). The T1w image is optimal for most cortical surface reconstruction pipelines (*e.g*., Freesurfer), while a microstructure-sensitive image (‘micro-image’) may be compiled in a variety of ways (e.g. R1 from MP2RAGE or MPM, or computation of T1w/T2w). The micro-image is then rigidly aligned to the surface-space T1w image. Finally, following generation of equivolumetric surfaces between the pial and GM/WM boundary, the intensities of the co-registered micro-image are sampled along the intracortical surfaces at matched vertices to produce microstructure profiles. **(Below)** 3D profile extraction is operationalised by sampling matched vertices on intracortical surfaces of various cortical depths. The protocol provides a consistent approach that can be used on different modalities, such as histology and MRI. Resulting profiles can be explored in a wide variety of ways including, computation of a matrix of regional similarities in microstructure (i.e. microstructure profile covariance) and gradients of microstructural differentiation (Paquola, Vos De Wael, *et al*., 2019); individual differences in microstructure (*e.g*., during development, (Paquola, Bethlehem, *et al*., 2019)) or fine-grained microstructural patterning with regions (Paquola *et al*., 2025).

The workflow is now available as a standalone toolbox (https://github.com/caseypaquola/cortpro) and incorporated within the open access multimodal MRI processing software “micapipe” (Cruces *et al*., 2022). The toolbox works flexibly on MRI and 3D histology, given a cortical surface reconstruction (pial and white matter boundary) and an aligned volume for sampling. To support integration with existing workflows and common imaging sequences, we have incorporated several optional preprocessing steps into the CortPro toolbox, including cortical surface reconstruction using FastSurfer (Henschel *et al*., 2020) and computation of a N3 bias-corrected T1w/T2w ratio image (Nerland *et al*., 2021). Therefore, the minimal requirements for the initiating the CortPro workflow are a T1w and T2w image (**Figure 1 top middle**).

Comparing MPs across areas, individuals or groups is simplified by using their central moments (**Figure 2A**). The zeroth moment (μ0) is the mean intensity and thus disregards variation in intensities across cortical depths. The higher moments (μ1-μ4) are, in contrast, sensitive to profile shape. The centre of gravity (μ1) indicates the balance of intensities in supra-vs-infragranular layers, while variance (μ2) reflects the spread of intensities across depths. For MRI-derived profiles, μ2 effectively indexes the flatness of the profile. Finally, skewness (μ3) and kurtosis (μ4) capture intensity shifts in the tails of the profiles (*i.e*., uppermost and lowermost depths) and for imaging data tend to be highly correlated with μ2 and μ1, respectively (|r| > 0.92, **Supplementary Figure 1**). Notably, the zeroth to second moments capture distinct patterns of microarchitectural variation (spatial correlation between moments: 0.34 < |r| < 0.60, **Figure 2A**). Therefore, moment maps offer complementary information on cortical differentiation. For *in-vivo* imaging, the central moments offer sufficient granularity and distinctiveness to describe the smooth profiles, but with higher resolution data more elaborate features may be appropriate.

**Figure 2.**
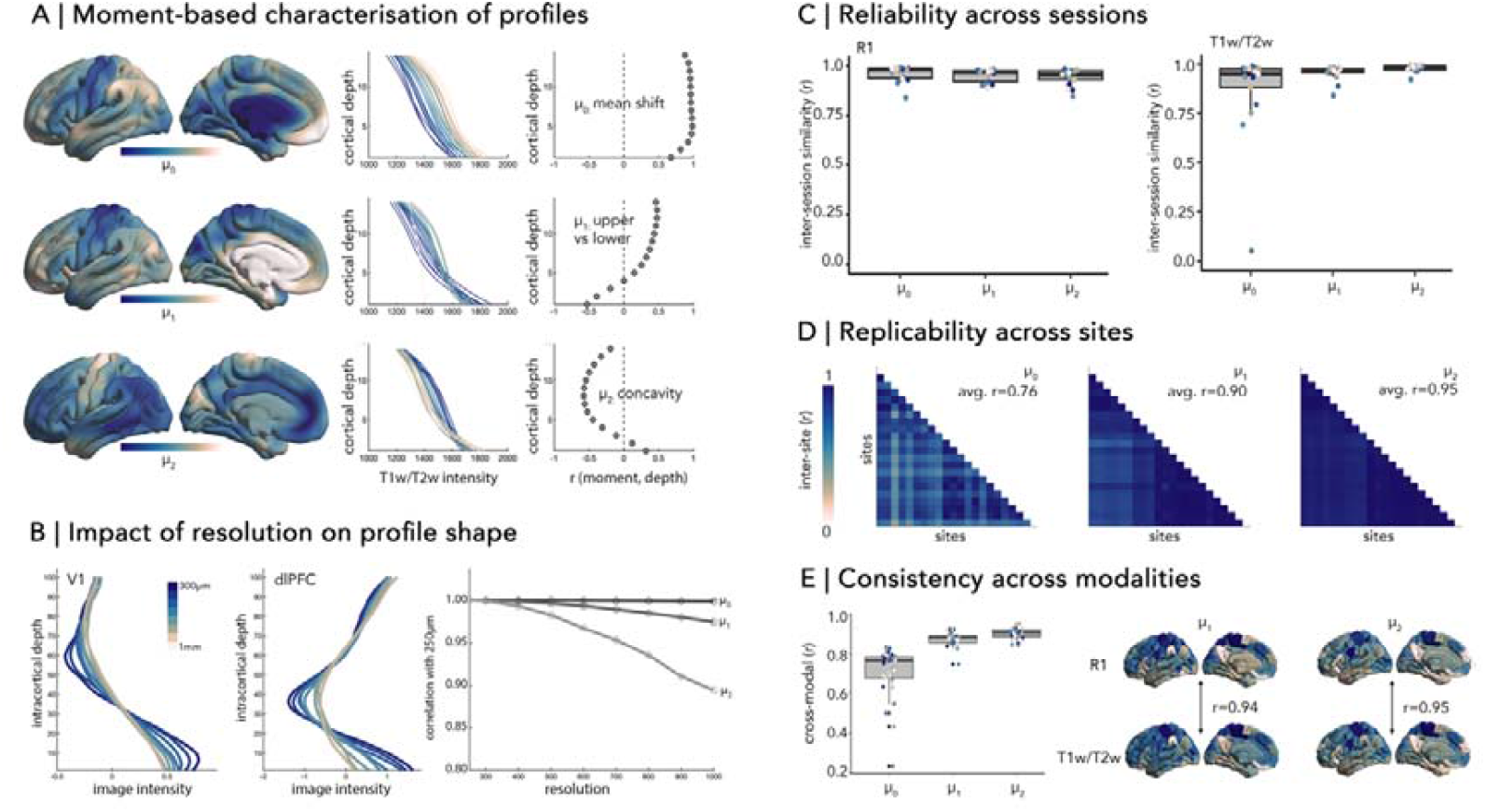
**A)** *(Left)* Regional variation in zeroth to second moment [R1 at 3T, (Royer *et al*., 2022)]. *(Middle)* Line plots represent average profiles within bins of the moment spectrum, with colours matched to the cortical plots, illustrating how the moments are related to change in profile shapes. *(Right)* Spatial correlation of moments with depth-wise intensities show differential sensitivities to shape. **B)** *(Left)* Intracortical profiles derived from increasingly downsampled structural MRI (original acquisition=0.25mm) acquired in one individual (Lüsebrink *et al*., 2021). *(Right)* Line plots show the similarity of cortex-wide moment maps computed on downsampled volumes, relative to the original ultra-high resolution volume (0.25mm). **C)** Boxplots show inter-session stability of profile moments, with points coloured by subject. **D)** Matrices illustrate the correlations between group-average moment maps derived from T1w/T2w imaging, acquired at 21 different sites. **E)** Boxplots show the within-subject correlations of moment maps derived from different modalities (R1 and T1w/T2w). Cortical surfaces depict group-average maps of μ1 and μ2 derived from R1 (*above*) and T1w/T2w (*below*) in the same sample. **Note:** Further details on these analyses can be found in the *Supplementary Information*.

### Proxying cortical lamination with MRI

Standard microstructure-sensitive imaging sequences are now able to reach sub-millimetre resolution with relatively short scan times but identifying cortical laminar or prominent myelin bands (*e.g*., stria of Gennari) depends upon even higher resolution imaging (*i.e*., < 0.5mm). Several studies have achieved this goal with *in-vivo* MRI, but they require long scan times or small fields of view that are typically infeasible to collect in large-scale or clinical studies. Therefore, an important assumption of the present approach is that the shape of the microstructure profile can still be resolved up to a certain degree at lower resolutions.

To test this point, we compared microstructure profiles and moment maps at a range of resolutions, based on the downsampling of a T1w image acquired at 0.25mm resolution in one individual (Lüsebrink *et al*., 2021). While the profiles are clearly smoother at lower resolutions, the moment maps are strikingly similar even at 1mm resolution (r > 0.9) (**Figure 2B**). At 0.7mm resolution, the correlations even exceed 0.95. Higher moments are increasingly sensitive to resolution (r_1mm_ μ0 = 1.0, μ1 = 0.98, μ2 = 0.90), which reinforces a selective focus on μ0-μ2 in *in-vivo* imaging studies. In sum, microstructure profiling with 0.7 - 1mm resolution imaging can still capture the regional organisation of cortical laminar.

### Reliability and replicability of MPs

As with any method, the utility of intracortical microstructure profiling in research and clinical studies is dependent upon reliability and replicability. To assess these attributes, we analysed several open-access datasets that involved multi-session, multi-modal imaging (Shams, Norris and Marques, 2019; Royer *et al*., 2022; Cabalo *et al*., 2025), or acquisition across sites with distinct scanning protocols (Casey *et al*., 2018). Comprehensive descriptions of these datasets and preprocessing are detailed in the *Supplementary Information*.

Test-retest (or scan-rescan) reliability provides insight into measurement error and is especially important for longitudinal studies that aim to capture meaningful intra-individual variation. To assess reliability, we compared individual-specific moment maps derived from different scanning sessions [n=17, 2 sessions within one day (Shams, Norris and Marques, 2019)]. The scanning protocols involved both quantitative T1 mapping (longitudinal relaxation time, “R1”) and non-quantitative acquisitions of T1w and T2w contrasts. Moment maps were very highly correlated across sessions for both R1 (r = 0.95 ± 0.04) and T1w/T2w (r = 0.95 ± 0.11) (**Figure 2C**), with the exception of μ0 in T1w/T2w imaging (r = 0.87 ± 0.23). This effect is likely related to B1+ transmit field biases that are known to influence T1w/T2w intensities (Glasser *et al*., 2022). Higher moments (μ1-μ4) sidestep this issue as their computation depends on relative values within an area, thus being relatively resilient to low spatial frequency variations that are characteristic of the B1+ field. Corroborating these results, we also found that the intraclass correlation coefficient (ICC), which is more sensitive to the discriminability of individuals, ranged from moderate to high across moments and regions, with the exception of low values for μ0 in T1w/T2w imaging [R1: ICC(3,1) = 0.70 ± 0.19, T1w/T2w: ICC(3,1) = 0.70 ± 0.30, **Supplementary Figure 2**]. Notably, these ICC values position the reliability of microstructure profiles on par with structural volumetry (Morey *et al*., 2010) and cortical thickness (Iscan *et al*., 2015) and superior to functional MRI (Elliott *et al*., 2020). We found similarly high levels of reliability in two additional datasets that involved longer durations between scans [PNI: n = 10, inter-session interval = 1.9 ± 2.5 months, inter-session similarity = 0.95 ± 0.04 (Cabalo *et al*., 2025). MICs: n = 40, inter-session interval = 1.71 ± 1.03 years, inter-session similarity = 0.89 ± 0.05 (Royer *et al*., 2022), **Supplementary Figure 2**]. Thus, the shape of microstructure profiles, as captured by μ1-μ4, is highly reliable, even if the reliability of the regional intensity values is not, as is the case for T1w/T2w.

Replicability reflects the independence of imaging features to site-specific effects. With global data aggregation necessary to grasp population diversity, ensuring the consistency of imaging features across sites is increasingly important. Thus, we examined the similarity of moment maps acquired across 21 sites, including 3 different scanner manufacturers [ABCD dataset; (Casey *et al*., 2018), **Supplementary Figure 3**]. Given the substantial sample size (n=5046) and the consistency of age distributions across sites (**Supplementary Figure 3**), we expected that any systematic differences in moment maps would primarily reflect scanner-related factors. Inter-site replicability of moment maps (product-moment correlation across sites) was very high, especially for μ2 (r = 0.95 ± 0.03) and μ1 (r = 0.90 ± 0.08) (**Figure 1D**). Replicability was lower for μ0 (r = 0.76 ± 0.13), which as above may be attributed to the influence of B1 transmit field bias. Together, these findings highlight the robustness of intracortical microstructure profiling in producing replicable metrics of cortical architecture that generalise across diverse scanner models and acquisition environments.

Finally, we evaluated whether intracortical microstructure profiles can be compared across modalities. Specifically, we assessed the similarity of moment maps derived from different myelin-sensitive contrasts (R1 and T1w/T2w) that were acquired within the same session for 17 individuals (Shams, Norris and Marques, 2019). At an individual level, cross-modal correlation of moment maps was very high for μ2 (r = 0.90 ± 0.03) and μ1 (r = 0.87 ± 0.03) (**Figure 1E**). In line with previous analyses, the consistency of μ0 was lower (r=0.71±0.10). We found a similar pattern of results with comparison of more distinct myelin-sensitive sequences [R1 and MTsat, (Cabalo *et al*., 2025)], whereby μ2 (r = 0.47 ± 0.12) and μ1 (r = 0.40 ± 0.10) were more similar across modalities than μ0 (r = 0.16 ± 0.13) (**Supplementary Figure 4**). Thus, examination of the shape of the profile can help to overcome idiosyncrasies of specific modalities and potentially bridge across datasets that use different sequences.

## Conclusion

With microstructure-sensitive imaging entering the mainstream of neuroimaging research, profiling offers a versatile and reliable approach for indexing cortical lamination. The present findings indicate that the shape of intracortical intensity profiles (as described by μ1 and μ2) offers a reliable and replicable basis for assessing inter-individual differences in cortical microstructure. Future studies would benefit from incorporating quantitative relaxometry (*e.g*., R□mapping) to facilitate region-wise comparisons of microstructural properties, as reliability and replicability decline when using non-quantitative contrasts such as T□w/T□w ratios. More broadly, our results suggest that 1 mm isotropic MRI is sufficient to capture meaningful laminar variation, though higher resolutions can provide measurable improvements in sensitivity and precision, and are becoming increasingly attainable with contemporary acquisition protocols (*e.g*., 0.65mm, MP2RAGE at 3T within ∼9 min (Bapst *et al*., 2024)).

To support future applications of the intracortical microstructure profiling approach, we developed a generalisable workflow for their extraction (https://github.com/caseypaquola/cortpro) and incorporated the code into an automated data preprocessing toolbox [micapipe, (Cruces *et al*., 2022)]. Furthermore, we’re releasing a warehouse of microstructure profiles and moment maps derived from open-access resources (“Microstructural Marketplace”, https://osf.io/e6f7d/). These include a wide range of microstructure-sensitive contrasts (R1, T1w/T2w, MTsat) acquired at various resolutions (0.25mm-0.8mm) and different magnetic strengths (3T and 7T) (Shams, Norris and Marques, 2019; Lüsebrink *et al*., 2021; Royer *et al*., 2022; Cabalo *et al*., 2025), as well as post-mortem staining for cells bodies [BigBrain: Amunts *et al*., 2013; Paquola *et al*., 2021)], myelin and parvalbumin interneurons [AHEAD: (Alkemade *et al*., 2022)]. We hope these datasets provide a helpful benchmark for new profiling studies and supports further investigations into the importance of microarchitecture in shaping brain function and cognition.

## Acknowledgements

We’d like to thank Dr. Kyesam Jung for beta testing CortPro.

This work was supported by the Deutsche Forschungsgemeinschaft (Emmy Noether Programme – 524408221).

BCB acknowledges research support from the National Science and Engineering Research Council of Canada (NSERC RGPIN-2025-05932), CIHR (FDN-154298, PJT-174995, PJT-191853, PJT-203761), SickKids Foundation (NI17-039), HIBALL, Healthy Brains and Healthy Lives (HBHL), Brain Canada Foundation, FRQS, the Tier-2 Canada Research Chairs Program, and the Centre for Excellence in Epilepsy at the Neuro (CEEN).

