## Supplementary for "Intracortical microstructure profiling: a versatile method for indexing cortical lamination"

**Supplementary Information**
In the following sections, we describe (1) data acquisition, (2) intracortical microstructure profiling and (3) statistical analyses of validity, reliability and replicability. For unrestricted datasets, the profiles and moments maps are made openly available in the "Microstructural Marketplace" (https://osf.io/e6f7d/).

### 1. Data acquisition
Key information on study size, demographics and imaging acquisition is provided in Supplementary Table 1. Detailed descriptions of the scanning protocols are provided below (directly quoted from the primary sources where possible).

**Supplementary Table 1: Dataset overview**

| *Dataset or first author* | *n* | *Age (years)* | *Male (%)* | *Modality* | *Resolution* | *Sessions (interval)* | *Included in OSF* |
|---|---|---|---|---|---|---|---|
| *Lüsebrink* | 1 | 27-38 | 100 | T1w | 0.25mm | 5 (up to 3mos) | Yes |
| *Shams* | 17 | 24.7 ± 2.8 | not reported | R1 and T1w/T2w | 0.8mm | 2 (same day) | Yes |
| *MICA-PNI* | 10 | 26.6 ± 4.60 | 40 | R1 and MTsat | 0.8mm | 2-3 (1.9±2.5mos) | Yes |
| *MICA-MICs* | 40 | 29.54 ± 5.62 | 54 | R1 | 0.8mm | 2 (1.71±1.03yrs) | Yes |
| *ABCD* | 5046 | 9.95 ± 0.62 | 51 | T1w/T2w | 1mm | 1 | No |

All studies reported that participants were healthy and provided written informed consent.

#### *1.1. Lüsebrink et al., 2021*
"All data were acquired on the same 7 T MR scanner (Siemens 7 T Classic, Siemens Healthineers, Erlangen, Germany). From 2009 to 2011 data were acquired with a 1 Tx/24 Rx channel head coil (Nova Medical Inc, MA, USA) and from 2012 onwards with a 1 Tx/32 Rx channel head coil (Nova Medical Inc, MA, USA). The scanner's software remained unchanged at VB17." Eight single T1-weighted MPRAGE volumes with prospective motion correction were acquired in five sessions across three months (MP-RAGE; 0.25 mm isotropic voxels, matrix = 880 × 880, 640 sagittal slices, TR = 3580 ms, TE = 2.41 ms, TI = 1210 ms, flip angle = 5°, bandwidth = 440 Hz/px, acquisition time ~ 53min). These volumes were averaged following non-linear registration to produce a 0.25mm T1-wighted image with high SNR.

#### *1.2. Shams, Norris and Marques, 2019*
"All scans were performed on a 3T Siemens Magnetom Prisma system (Siemens, Erlangen, Germany) equipped with a 32-channel receive coil."

"All the following experiments were run twice for each subject, test and retest, with the subject being repositioned in the scanner in between the two sessions. The subjects' heads were either simply held with head cushions or additionally fixed using a chin-rest (to avoid head translation and rotation in the head to foot direction) in the test and retest condition (the fixation was randomized). Also, in both the test and retest condition, the order of the T1w/T2w imaging protocols and R1 mapping protocols were randomized."

"MP2RAGE sequence parameters were: $TR/TI_1/TI_2$ = 5.5/0.7/2.5 s, BW = 210 Hz/Px, Flip angles ($\alpha1/\alpha2$) = 6°/4°, parallel imaging = 3, TE/echo spacing = 3.28/7.5 (ms/ms), FOV = 256 × 256 × 204.8, matrix size = 320 × 320 × 256, Partial Fourier in slice encoding direction = 6/8 (resulting in 64 and 128 excitations before and after the k-space center), Tacq = 12min 30sec. To attain more accurate cortical surfaces, we acquired high spatial resolution images of 0.8 × 0.8 × 0.8 $mm^3$."

"The fast saturation recovery turbo FLASH acquisition had the following parameters: TR/TE = 10/2.23 (s/ms), BW = 490 Hz/Px, FA = 8°, spatial resolution = 3.3 × 3.3 × 2.5 $mm^3$, Dist. Factor = 100%, Tacq = 20 sec."

"T1w and T2w scans were collected using a T1w magnetization-prepared rapid gradient echo (MPRAGE) and a T2w sampling perfection with application optimized contrast using different angle evolutions (SPACE) sequences adapted from the Human Connectome Project imaging protocol. The 3D MPRAGE images were acquired with the parameters of: TR = 2200 ms, TE = 2.64 ms, TI = 1100 ms, 11° flip angle, bandwidth = 170 Hz/pixel, echo spacing = 9.3 ms, FOV 256 mm × 320 mm × 179.2 mm, matrix 320 × 400 × 224, 0.8 mm isotropic resolution, iPAT = 3, and an acquisition time of 6 min and 2 sec. The acquired SPACE images had the following parameters: TR = 3200 ms, TE = 569 ms, variable flip angle optimized for T2 weighting, bandwidth = 579 Hz/pixel, echo spacing = 4.12 ms, Turbo Factor = 314, FOV 256 mm × 320 mm × 179.2 mm, matrix 320 × 400 × 224, 0.8 mm isotropic resolution, iPAT = 3 and acquisition time of 6 min and 19 sec. T1w and T2w images were receive bias field corrected as provided by the vendor based on additional reference volume coil scan (Siemen's pre-scan normalize)."

"B0 field maps were acquired for the purpose of correcting readout distortion in the T1w, T2w and R1 images using a dual-echo gradient echo sequence adapted from the Human Connectome Project imaging protocol (http://humanconnectome.org/study/hcp-young-adult/documentation/data-release/Q1_Release_Appendix_I.pdf) with delta TE = 2.46 ms. Other imaging parameters are as follows: TR = 731 ms, TE1 = 4.96 ms, 50° flip angle, bandwidth = 566 Hz/pixel, FOV = 104 mm × 90 mm × 72 mm, matrix = 208 × 180 × 144, 2 mm isotropic resolution, interleaved multi-slice mode and the acquisition time of 2 min and 15 sec."

*1.3. MICA-PNI (Cabalo et al., 2025)*

"MRI data were acquired on a 7 T Terra Siemens with a 32-receive and 8-transmit channel head coil in parallel transmission (pTX) mode." Across three sessions, "3D-magnetization-prepared 2-rapid gradient-echo sequence with Universal Pulses to optimize $B1^+$ uniformity were acquired (MP2RAGE; 0.5 mm isovoxels, matrix = 320 × 320, 320 sagittal slices, TR = 5170 ms, TE = 2.44 ms, TI1 = 1000 ms, TI2 = 3200 ms, flip = 4°, iPAT = 3, partial Fourier = 6/8, FOV = 260 × 260 $mm^2$)". In the third session, "one myelin-sensitive magnetization transfer (MT; 0.7 mm isovoxels, TR = 95 ms, TE = 3.8 ms, flip angle = 5°, FOV = 230 × 230 $mm^2$, slice thickness = 0.72 mm, 240 sagittal slices)" was acquired.

*1.4. MICA-MICs (Royer et al., 2022)*

"Scans were completed at the Brain Imaging Centre of the Montreal Neurological Institute and Hospital on a 3 T Siemens Magnetom Prisma-Fit equipped with a 64-channel head coil. Two T1w scans with identical parameters were acquired with a 3D magnetization-prepared rapid gradient-echo sequence (MP-RAGE; 0.8 mm isotropic voxels, matrix = 320 × 320, 224 sagittal slices, TR = 2300 ms, TE = 3.14 ms, TI = 900 ms, flip angle = 9°, iPAT = 2, partial Fourier = 6/8). Both T1w scans were visually inspected to ensure minimal head motion before they were submitted to further processing. qT1 relaxometry data were acquired using a 3D-MP2RAGE sequence (0.8 mm isotropic voxels, 240 sagittal slices, TR = 5000 ms, TE = 2.9 ms, TI 1 = 940 ms, T1 2 = 2830 ms, flip angle 1 = 4°, flip angle 2 = 5°, iPAT = 3, bandwidth = 270 Hz/px, echo spacing = 7.2 ms, partial Fourier = 6/8). We combined two inversion images for qT1 mapping in order to minimise sensitivity to B1 inhomogeneities and optimize intra- and inter-subject reliability".

*1.5 Adolescent Brain Cognitive Development (ABCD) (Casey et al., 2018)*

"The ABCD imaging protocol is harmonized for three 3T scanner platforms (Siemens Prisma, General Electric (GE) 750 and Philips) and use of multi-channel coils capable of multiband echo planar imaging (EPI) acquisitions, using a standard adult-size coil." Each scanning session involved acquisition of a 3D T1-weighted magnetization-prepared rapid acquisition gradient echo (MP-RAGE) scan and 3D T2-weighted fast spin echo (FSE) with variable flip angle scan. For both sequences and across all sites, scans were acquired with an isotropic voxel resolution of 1mm and FOV = 256 x 256. Parameters specific to the scanner manufacturer are detailed in Supplementary Table 2.

**Supplementary Table 2: ABCD imaging scanning parameters**

| Scanner | Sequence | Slices | TR (ms) | TE (ms) | TI (ms) | Flip Angle (deg) | Parallel Imaging | Acquisition Time |
|---|---|---|---|---|---|---|---|---|
| Siemens (Prisma VE11B-C) | T1 | 176 | 2500 | 2.88 | 1060 | 8 | 2× | 07:12 |
| | T2 | 176 | 3200 | 565 | N/A | Variable | 2× | 06:35 |
| Philips (Achieva dStream, Ingenia) | T1 | 225 | 6.31 | 2.9 | 1060 | 8 | 1.5 × 2.2 | 05:38 |
| | T2 | 256 | 2500 | 251.6 | N/A | 90 | 1.5 × 2.0 | 02:53 |
| GE (MR750, DV25-26) | T1 | 208 | 2500 | 2 | 1060 | 8 | 2× | 06:09 |
| | T2 | 208 | 3200 | 60 | N/A | Variable | 2× | 05:50 |

For the present study, we used raw baseline imaging data from the 5th release of ABCD. Inclusion criteria were T1w and T2w images that passed quality control (Hagler *et al.*, 2019), as well as successful completion of Freesurfer (run in-house, v7.2.0), resulting in 5046 individuals.

### 2. Intracortical microstructure profiling

This section outlines the intracortical microstructure profiling procedure using "CortPro" (https://github.com/caseypaquola/CortPro).

*A) Surface Reconstruction*

Pial and white matter (GM/WM boundary) cortical surfaces are reconstructed from native T1-weighted (T1w) images using FastSurfer (Henschel *et al.*, 2020). If multiple T1w scans are available for a subject, they are first rigidly aligned and averaged to improve signal-to-noise ratio before surface reconstruction. Alternatively, users may supply pre-existing surfaces, provided they conform to the FreeSurfer/FastSurfer format.

Following surface reconstruction, a series of intracortical surfaces are generated between the pial and GM/WM surfaces using an equivolumetric layering approach (Waehnert *et al.*, 2014; Wagstyl *et al.*, 2018). By default, 14 intracortical surfaces are created, though this number can be modified via an optional command-line argument.

*B) Volume Processing*

Microstructure-sensitive images are produced in various ways depending on the acquisition sequence. Given the widespread availability of T1w and T2w scans, the toolbox supports the computation of T1w/T2w ratio images.

If multiple T1w or T2w acquisitions are available, they are rigidly aligned and averaged within each modality. Intensity nonuniformities are then corrected using N3 bias correction (Sled, Zijdenbos and Evans, 1998; Nerland *et al.*, 2021). The bias-corrected T2w image is then rigidly aligned to the bias-corrected T1w image using ANTs (Avants *et al.*, 2008), and the voxelwise T1w/T2w contrast is computed.

If surface reconstruction and T1w/T2w images are generated within the toolbox, the T1w/T2w volume is resliced into surface space using FreeSurfer's mri_vol2vol with the --regheader flag. If users provide external surfaces or a precomputed microstructure-sensitive image, these inputs are rigidly co-registered via the following process:

i) Brain segmentation is performed on both the T1w structural image (orig.mgz) and the microstructure-sensitive image using SynthSeg (Billot *et al.*, 2023),
ii) An affine registration matrix is computed between the two brain-extracted images using ANTs (Avants *et al.*, 2008)
iii) This transform is then applied to the full-head microstructure-sensitive image to bring it into alignment with the surface space.

*C) Microstructure profiling*

The co-registered microstructure-sensitive image is then sampled across intracortical surfaces (Freesurfer 7.4, mri_vol2surf). As the intracortical surfaces preserve the vertex indexing of the pial and white matter surfaces, sampled intensity values across depths can be organised into a matrix of depths x vertices, with columns representing vertex-wise microstructure profiles.

*D) Shape analysis of profiles*

The shape of each intracortical microstructure profile is characterised by its central moments. The zeroth moment ($\mu 0$) is calculated as the mean intensity of the profile. Computation of the higher moments ($\mu 1$-$\mu 4$) involves treating the profile as a histogram with intracortical depths as bins and the intensities as frequency. To do so, we generate a data distribution that, when in histogram form, captures the profile shape. Then, we calculate the mean ($\mu 1$), standard deviation ($\mu 2$), skewness ($\mu 3$) and kurtosis ($\mu 4$) of the reformatted data. For *in vivo* microstructure profiles, only $\mu 0$-$\mu 2$ provide unique information (**Supplementary Figure 1**), though higher moments can provide more comprehensive characterisation in higher resolution datasets with more sampling depths.

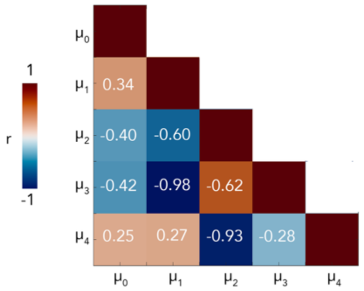


**Supplementary Figure 1:** Spatial (product-moment) correlations between group-average moment maps derived from R1 (MICA-MICs), showing uniqueness across $\mu 0$-$\mu 2$ ($0.34<|r|<0.60$), but redundancy of $\mu 1$ with $\mu 3$ ($|r|=0.98$), and $\mu 2$ with $\mu 4$ ($|r|=0.93$).

As described above, the toolbox is compatible with both raw and partially preprocessed data. For the current study, datasets were used in their most advanced stage of relevant preprocessing, as detailed in **Supplementary Table 3.**

**Supplementary Table 3: Level of preprocessing completed prior to toolbox implementation**

| *Dataset/first author* | *Preprocessed surfaces* | *Preprocessed volumes* |
|---|---|---|
| *Lüsebrink* | Freesurfer output provided | - |
| *Shams* | - | Bias corrected R1, T1w and T2w |
| *MICA-PNI* | Fastsurfer output provided | MTsat |
| *MICA-MICs* | Freesurfer output provided | - |
| *ABCD* | Freesurfer output provided | - |

### 3. Analyses of validity, reliability and reproducibility

<u>*Sampling experiment*</u>

To identify an appropriate sampling density for *in vivo* microstructure profiling, we counted the number of unique values within vertex-wise microstructure profiles from a healthy adult (MICA-MICs). The profiles were generated with 100 sampling points within the cortex, which we expected to vastly exceed the maximum unique values per profile. We repeated the procedure with nearest neighbour and trilinear interpolation. The number of unique values is correlated with cortical thickness in the case of nearest neighbour interpolation ($r=0.83$, $p<0.001$), but this effect is diminished with trilinear interpolation ($r=0.42$, $p<0.001$), further supporting the adoption of the latter. Nevertheless, the nearest neighbour approach provides a conservative benchmark for sampling density. With this individual, the maximum voxels directly along a profile was 15. Additionally, the mid-point between the number of unique values in nearest neighbour and trilinear approaches was 20±6. We previously established a standard sampling density of 14 for *in vivo* microstructure profiling, because at this sampling density the correlations between microstructure profiles reach peak stability (Paquola, Vos De Wael, *et al.*, 2019). Thus, given 14 is near the conservative benchmark and the more liberal mid-point between the two approaches, we continue to use it as a default sampling density for *in vivo* microstructure profiling.

<u>*Resolution experiment (Dataset: Lüsebrink et al., 2021)*</u>

We downsampled a 0.25mm T1w volume to a range of lower resolutions (0.3-1mm in steps of 0.1) using imresize3 in MATLAB. Then, we constructed microstructure profiles on the original and downsampled volumes using 100 intracortical depths and computed moment maps. Finally, we correlated the moment maps derived from downsampled volumes with the moment maps from the original volume to evaluate the effect of voxel resolution on regional variability in microstructure profile shapes.

*Reliability experiment (Datasets: Shams et al., MICA-MICs, MICA-PNI)*

For each participant, we calculated the product-moment correlation between moment maps derived from different scanning sessions to evaluating the stability of regional variations in profile shapes. Additionally, we calculated the intraclass correlation coefficient for each region and each moment as a measure of their internal consistency. For the reliability experiments, microstructure profiles were averaged within 200 spatially-contiguous parcels prior to moment calculation.

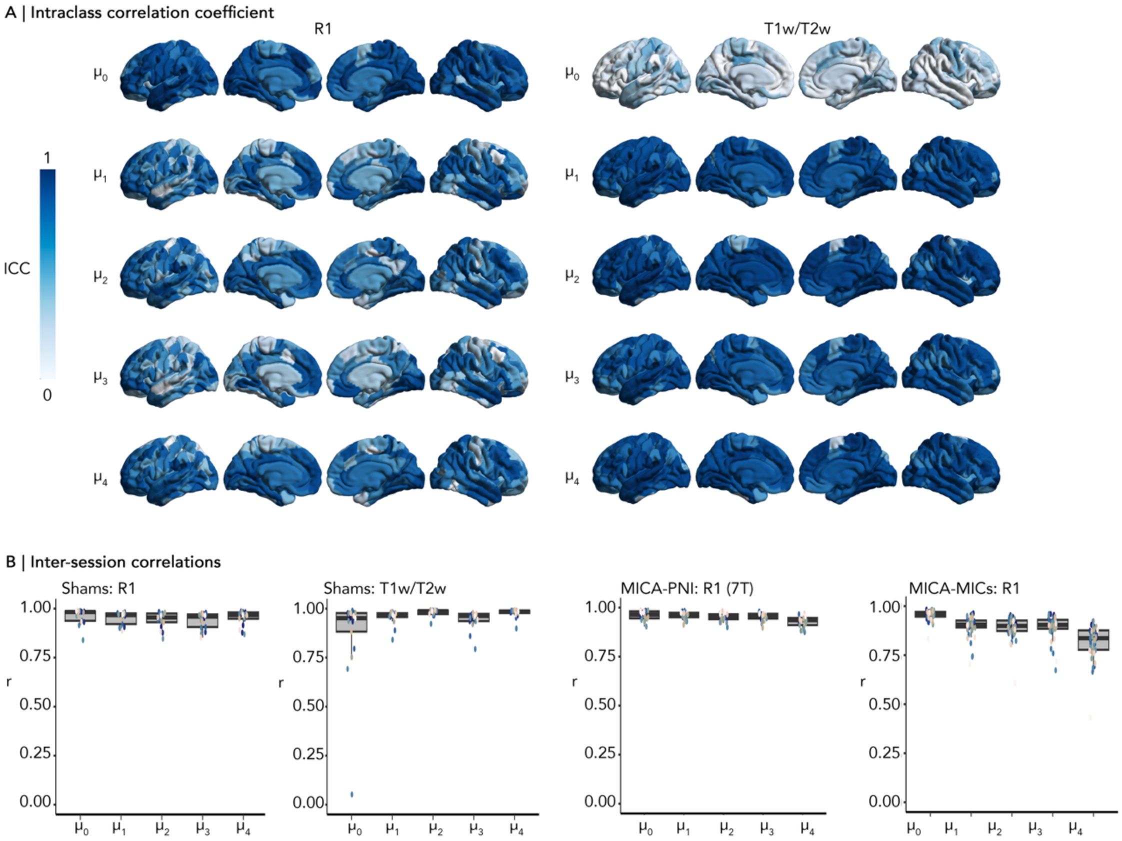


**Supplementary Figure 2:** A) Region-wise intraclass correlation coefficient for both R1 and T1w/T2w (Shams dataset, n=17, 2 sessions per individual). B) Boxplots depict the range of inter-session correlations across individuals in each dataset.

*Replicability experiment across sites (Dataset: ABCD)*

For each site of the ABCD study, we computed group-average moment maps on parcellated microstructure profiles. Then, we calculated the product-moment correlation of moment maps between each pair of sites. The age distributions were similar across sites (**Supplementary Figure 3).**

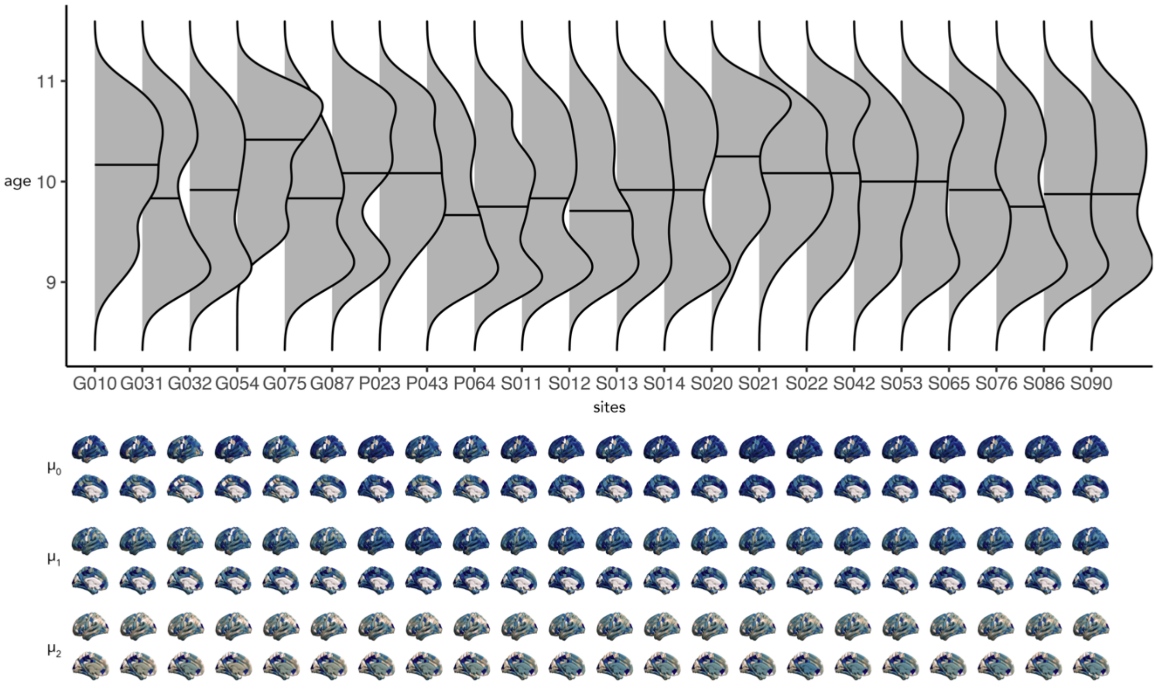


**Supplementary Figure 3:** (Above) Ridgeplots show the similar distribution of participant ages at each site for data included from the ABCD study. (Below) Site-average moment maps.

*Consistency experiment across modalities (Datasets: Shams et al., MICA-PNI)*

For each participant, we calculated the product-moment correlation of moment maps derived from different modalities within the same session. For the Shams dataset, this involved R1 and T1w/T2w in two sessions per individual, while in MICA-PNI this involved R1 and MTsat in one session per individual. The cross-modal correlations illustrate the generalisability of the approach across acquisition sequences, while also indicating the degree to which different "myelin-sensitive" protocols capture similar patterns of regional variation (**Supplementary Figure 4**).

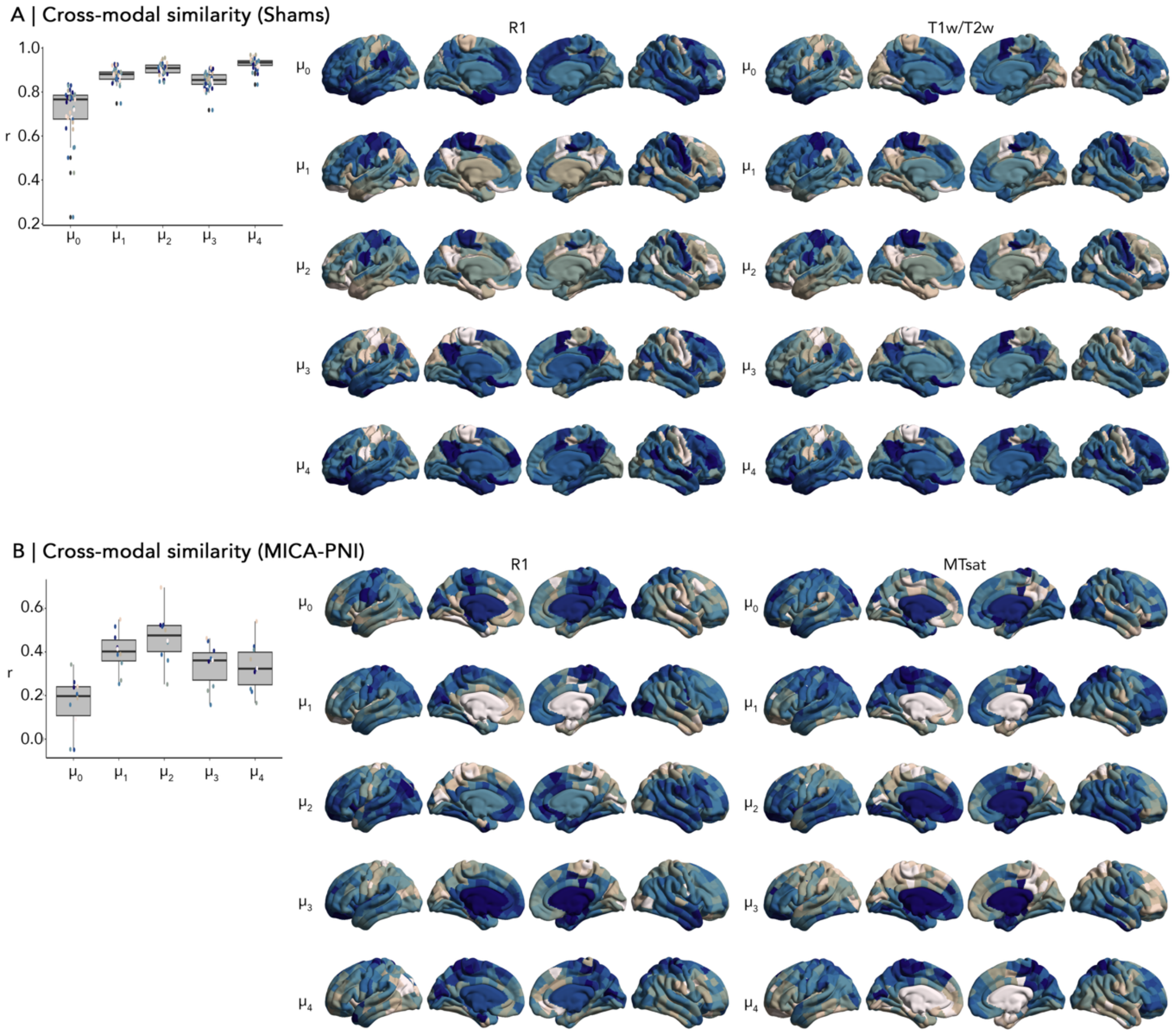


**Supplementary Figure 4:** Boxplots show the range of cross-modal correlations across individuals for each moment. Cortical plots show group-average moment maps for the various "myelin-sensitive" modalities.
